# Signaling pathway evaluation of leading ATRi, PARPi and CDK7i cancer compounds targeting the DNA Damage Response using Causal Inference

**DOI:** 10.1101/2024.02.15.580418

**Authors:** Nina Truter, Wikus Bergh, Martine van den Heever, Shade Horn, Klarissa Shaw, Dan Leggate, Neil Wilkie, Raminderpal Singh

## Abstract

**Introduction:** There are many cancer drugs in development which target the DNA damage response (DDR), following early successes of drugs such as olaparib. However, various challenges to the success of these inhibitors exist, including the emergence of resistance, the identification of appropriate biomarkers to identify patients who will respond to treatment, as well as the identification of combination therapies that improve efficacy without a concomitant increase in toxicity. While the identification of biomarkers of resistance could aid in overcoming these challenges, current methods mostly generate lists of potential genes, proteins that display changes in cancer patients, without exposing the underlying, and often critical, mechanisms of resistance.

**Methods:** We have developed the Adaptable Large-Scale Causal Analysis (ALaSCA) software platform, which applies Pearlian Causal Inference (PCI) techniques to specifically transcriptomic, proteomic and phenotypic multi-omics data. ALaSCA quantifies the causal contributions of different biological pathways to an outcome such as responsiveness to treatment. The strength of applying PCI to biological pathways lies in quantifying the causal contributions of targets, through their related pathways, to drug sensitivity - as opposed to merely enriching or grouping lists of genes into pathways. We applied ALaSCA to transcriptomic data for several different compounds related to three known inhibitor types that target DDR proteins: an ATR, a CDK7, and several PARP inhibitors. Our aim was to use causal methods to evaluate biological signaling pathways to identify resistance mechanisms that can be used for patient stratification and development of combination therapies in breast, ovarian and non-small cell lung cancer (NSCLC).

**Key findings:** We observed that niraparib seems to have a different resistance mechanism than other PARPi inhibitors in breast and NSCLC, which is driven by CDK1 as opposed to base excision or nucleotide excision repair. Additionally, CDK7 appears to be a significant driver of PARP inhibitor resistance, especially for niraparib, through predominantly the G2/M cell cycle phase and to a lesser extent nucleotide excision repair in breast and ovarian cancer, but not in NSCLC. Lastly, we identified that genes from the homologous recombination pathway drive resistance to AZD6738, an ATR inhibitor, in breast cancer, and that AZD6738 and irinotecan have differing resistance mechanisms in ovarian cancer, indicating the potential of combining these treatments.

**Next steps:** Our findings demonstrate the potential of ALaSCA to generate interesting insights for treatments when applied to public data and well-known inhibitors. Partnership with industry drug discovery groups using proprietary data to rerun the above evaluations will further refine and confirm these findings.

## 1. Introduction

There are many cancer drugs in development which target the DNA damage response (DDR), following early successes of drugs such as olaparib. There are several challenges posed to the success of these inhibitors, including emergence of resistance, identification of biomarkers for patient selection and identification of combination therapies that improve efficacy without a concomitant increase in toxicity (Chen *et al*., 2022). After the approval of PARP inhibitors (PARPi) and establishment of their clinical use, for example, resistance mechanisms such as reversion of the homologous recombination phenotype were uncovered (Lord and Ashworth, 2017; Baxter *et al*., 2022). There are, however, many patients displaying PARPi resistance for unknown reasons (Baxter *et al*., 2022). It is therefore likely that emerging therapies such as ATR and CDK7 inhibitors will face similar challenges as they make their way into the clinic. Indeed, preclinical studies have identified the role of *Cyclin E, CDK2* or *Myc* mutation or loss in ATR inhibitor (ATRi) resistance (Baxter *et al*., 2022). The identification of biomarkers of resistance aids in overcoming these challenges, however many methods result in lists of potential genes/proteins or pathways that display changes in cancer patients that do not take the mechanism of resistance into account.

We have developed the Adaptable Large-Scale Causal Analysis (ALaSCA) software platform, which applies Pearlian Causal Inference (PCI) techniques to multi-omics data, specifically transcriptomic, proteomic and phenotypic data (Louw *et al*., 2023) to quantify the causal contributions of different biological pathways to a phenotype such as treatment resistance.

The strength of applying PCI to biological pathways lies in quantifying the causal contributions of genes through their related pathways to drug sensitivity. The method creates a network diagram with weighted relationships between genes and relevant sections of their related pathways that drive drug sensitivity. These findings therefore produce more than just lists of individual gene targets, by contextualizing how the interconnection of signaling pathways contributes towards drug sensitivity. This also differs from analyses such as differential pathway expression where change in a pathway is used as an indicator of involvement in an outcome such as responsiveness to treatment. PCI does not only consider the change of genes from a baseline state, but rather the change of genes in relation to each other and in relation to the outcome.

ALaSCA enables scientists to evaluate biological signaling pathways through causal methods to identify potential resistance mechanisms that can be used for patient stratification and development of combination therapies. We applied ALaSCA to several compounds related to three inhibitors that target DDR proteins - an ATR inhibitor, CDK7 inhibitor (CDK7i) and several PARPis - to identify such opportunities in and around these compound families.

## 2. Methods

We applied ALaSCA to multi-omics data from GDSC, Cancer Cell Line Encyclopedia (CCLE) and Genentech to quantify the causal contribution of five DDR pathways to ATRi, CDK7i and PARPi resistance. Here, we used the IC50 values for each inhibitor in breast, ovarian, and non-small cell lung cancer (NSCLC) cell lines. Additionally, we applied ALaSCA to chemotherapies commonly used in breast and ovarian cancer, such as irinotecan and gemcitabine and separately noted the contribution of the ATR, CDK7 and PARP1/2 pathways to drug sensitivity of ATRi, CDK7i and PARPi, to identify their role in each other’s resistance mechanisms. The findings presented in section 3 include observations that can significantly impact the use of the three main DDR inhibitors.

### 2.1. Input data

ALaSCA was applied to RNASeq data from Garcia-Alonso *et al*. (2017). This consists of data for 1362 cell lines with mRNA expression for 15,379 genes (merged from GDSC, CCLE and Genentech), that is pre-processed, normalized, batch-corrected, and filtered to remove low expressed genes and samples by the original authors. The data are available at: https://www.synapse.org/#!Synapse:syn10463688/wiki/463140, accessed on 22 June 2023.

The relevant drug IC50s were obtained from the GDSC database, using IC50s from GDSC2 when available in preference to GDSC1 (https://www.cancerrxgene.org/), accessed on 29 November 2023. We used AZD6738 as an example of ATR inhibitors, THZ21021 as an example of CDK7 inhibitors and niraparib, veliparib, talazoparib, rucaparib and olaparib to compare findings between PARP inhibitors. We also included irinotecan and gemcitabine in our model.

### 2.2. ALaSCA platform

The ALaSCA platform is built using Python 3 and open-source python libraries, such as Numpy, Pandas, and Scipy, and PyWhy (DoWhy) (https://github.com/py-why/). The causal theory behind the ALaSCA simulation platform and how it was developed is discussed in more detail in the publication by Louw *et al*. (2023). In summary, PCI is conducted with the use of directed acyclic graphs (DAGs) which are the visual representation of the cause-and-effect diagram of the biological components in a mechanism, in our case proteins, genes, phenotypes and other similar features such as drug resistance (Figure 1).

**Figure 1.**
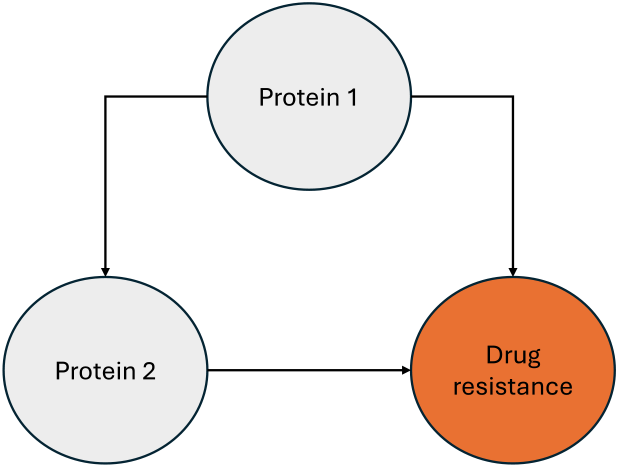
A simple example DAG showing the directed effects of two proteins in a pathway that drive drug resistance.

The biological components are represented by nodes and are connected by directed edges, which represent the causal relationships between the components. The DAG structure is based on prior knowledge of the biological or disease mechanism and key interactions between proteins within pathways of interest identified in literature. The strength of the causal effects, as represented in the DAG, are quantified during causal inference with the use of PCI (Glymour, Pearl and Jewell, 2016; Pearl, 2000). The ALaSCA platform can quantify these causal effects using a wide range of data types, such as clinical and other observational data, omics-type study data, as well as combinations thereof (Louw *et al*., 2023; Truter *et al*., 2022). The output that is generated in this study is represented by a causal value (“relative causal contribution” in our results) for each gene in the DAG, which indicates how strongly the downstream pathways of that gene are predicted to drive changes in the outcome such as drug IC50 values. Positive values indicate that the gene’s downstream pathways contribute to innate resistance to the drug (higher IC50 values) and negative values indicate that the gene’s downstream pathways contribute to innate sensitivity to the drug (lower IC50 values).

### 2.3. DAG construction

The DAG which was created for this study included five pathways involved in the DDR - base excision repair (BER), nucleotide excision repair (NER), homologous recombination (HR) and the G1/S and G2/M cell cycle phases (Figure 2, high quality version available in Supplemental Material). These pathways were chosen because our DDR inhibitors target a specific protein in each of these pathways and resistance often involves compensation in one of these pathways. For example, PARPi resistance can develop due to restoration of the HR pathway (Chen *et al*., 2022). This also enables the detection of insights of where the protein target is involved in the resistance mechanism of other compounds. We used pathway databases and relevant DDR literature to construct our causal diagram - each of the five pathways are directed to affect a compound’s sensitivity, represented by its IC50 in each cell line. We test the robustness of our constructed DAGs using methodology described previously (Louw *et al*., 2023).

**Figure 2.**
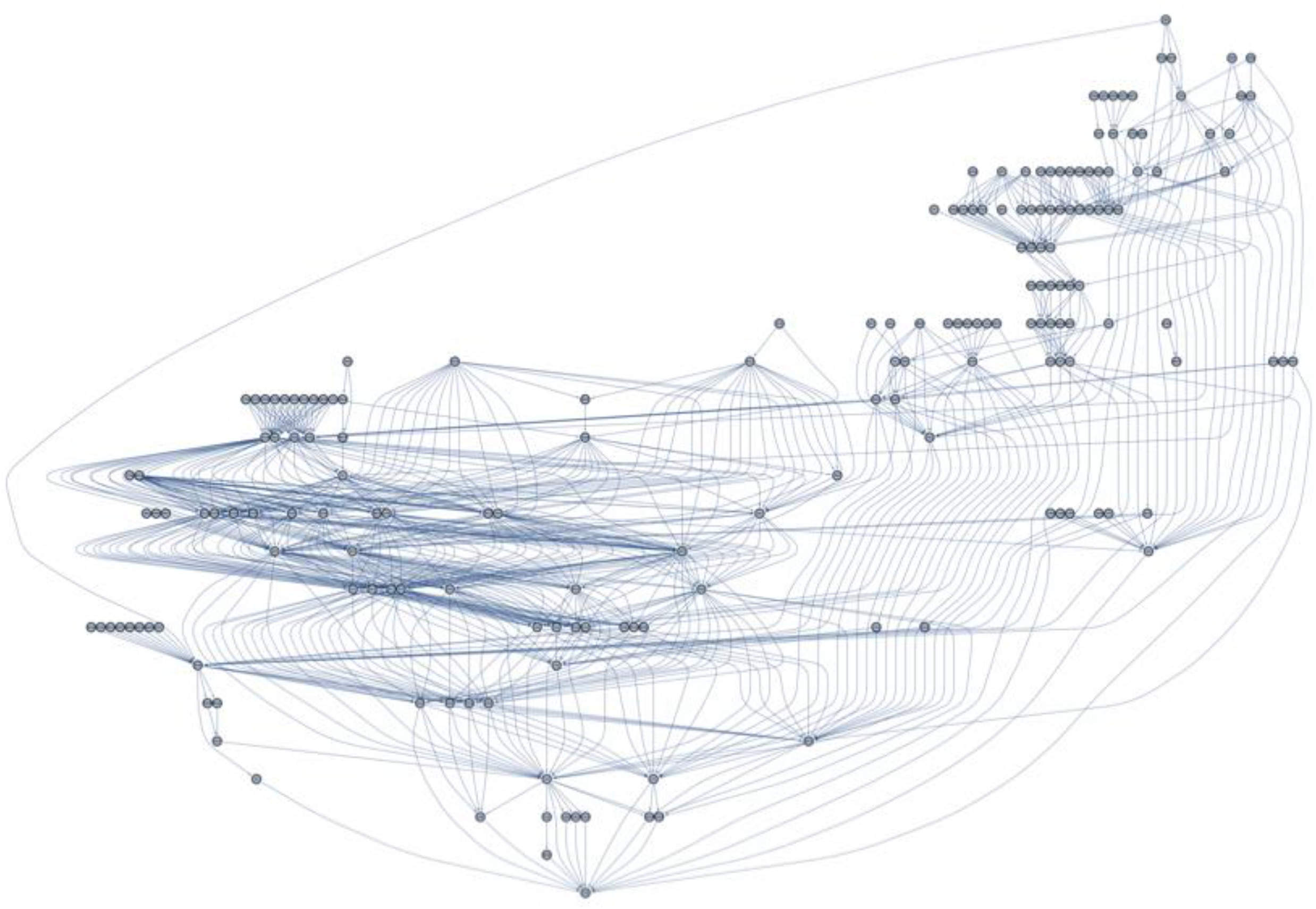
DAG containing genes involved in the BER, NER, HR pathways and the G1/S and G2/M cell cycle phases and how they are hypothesized to affect drug sensitivity, represented by IC50 values.

## 3. Results

We present the results separately based on the impact that the findings can have on each specific inhibitor. Firstly, by looking at the drivers of resistance for different PARPi across the cancers, secondly the opportunity for CDK7i due to the involvement of CDK7 in PARPi resistance and separately drivers of CDK7i resistance. Lastly we investigate ATRi resistance and how it differs from two chemotherapies in ovarian cancer. The relative causal contributions can be compared between inhibitors and cancers.

### 3.1. Signaling pathway evaluation of PARP inhibitors such as niraparib, veliparib, talazoparib, rucaparib and olaparib

Overall, for most PARPi across the cancers analyzed, we observed the involvement of G1/S and G2/M cell cycle genes. As a key example, the top 10 genes driving Niraparib resistance in NSCLC include several G2/M (WEE1, PLK1, PKMYT1, CKS2, CHEK1, Cyclin B2 and B3) and some G1/S cell cycle genes (CDC25A, E2F2, E2F3) (Figure 3). Our results indicate that CDK1 is the pivotal point for all our tested PARPi in ovarian cancer due to all the upstream pathways that causally contribute to resistance through it. This is supported by the observation that CDK1 depletion after paclitaxel treatment in combination with PARP inhibition significantly reduces cell growth of HR-proficient ovarian cancer cells (Yanaihara *et al*., 2023). Here a pivotal point refers to a gene that is involved in several of the pathways from the top 10 genes that drive PARPi resistance - for example, CDK1 is downstream of all seven G2/M cell cycle genes driving Niraparib resistance in NSCLC (Figure 3). Some of these genes have more than one downstream pathway that is also involved in resistance (e.g. CHEK1 affects both WEE1 and CDC25A, which both also contribute to resistance through CDK1). Table 1 lists the number of times CDK1 is downstream from the several pathways of these driving genes.

**Table 1:** the number of times CDK1 is involved in the pathway of the top 10 genes contributing to PARPi resistance in ovarian, breast and NSCL cancer.

|  | Niraparib | Olaparib | Rucaparib | Veliparib | Talazoparib |
| --- | --- | --- | --- | --- | --- |
| Ovarian | 12 | 10 | 11 | 12 | 10 |
| Breast | 5 | 4 | 3 | 13 | 2 |
| NSCLC | 12 | 6 | 0 | 1 | 0 |

**Figure 3.**
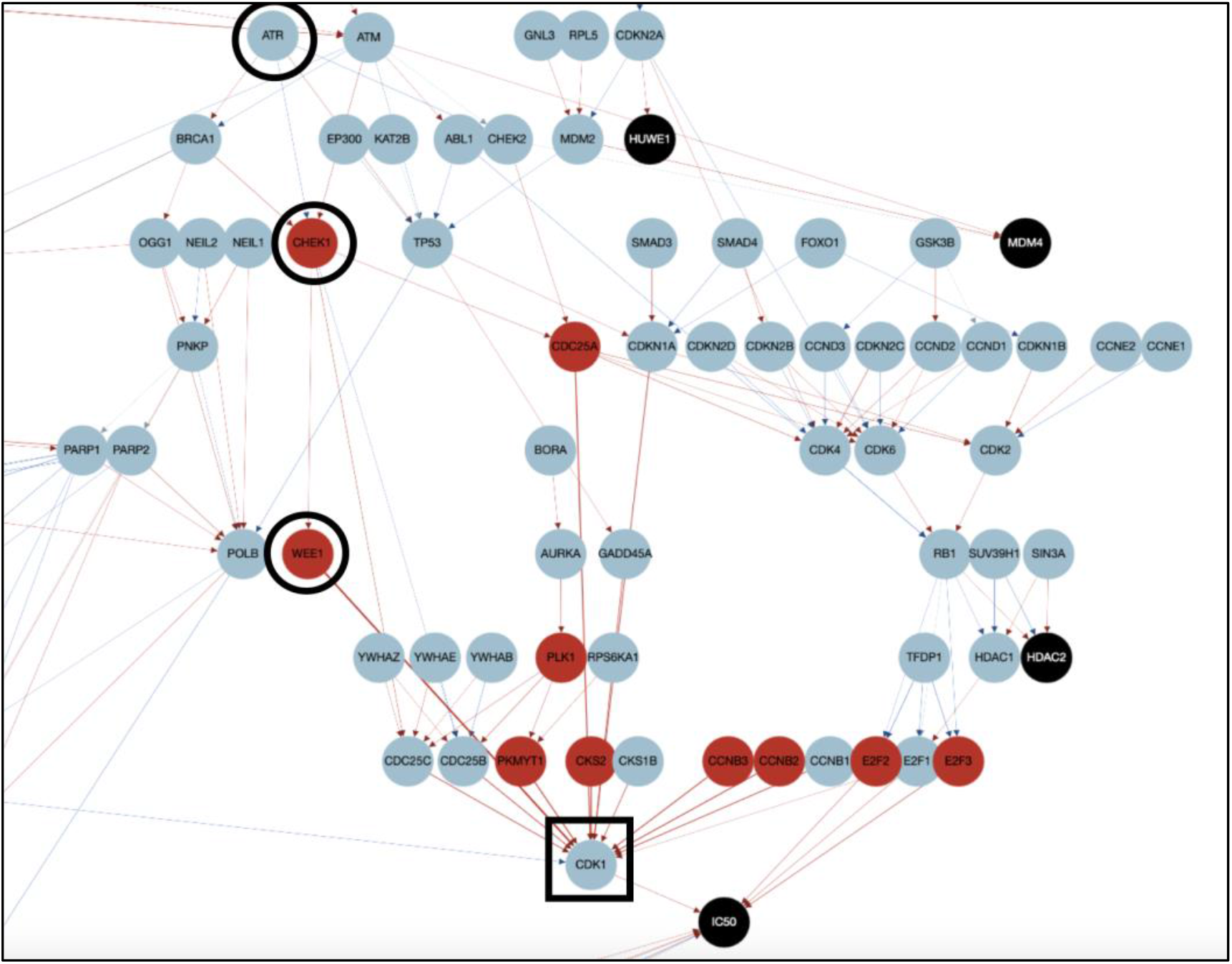
Portion of DAG indicating possible DDR pathways driving niraparib resistance in NSCLC. Red circles = genes upstream of CDK1 that drive resistance; black square = CDK1; black circled genes = ATR, CHEK1, WEE1.

However, in our results CDK1 only remains a pivotal point for veliparib in breast cancer and niraparib in NSCLC (Table 1). Additionally, CDK1’s causal contribution to PARPi resistance remains the strongest for niraparib and veliparib compared to the other PARPi’s across all three cancers (Figure 4). This may mean that there is a unique opportunity for these specific PARPi’s in NSCLC and breast cancer, either by using CDK1 as a predictive biomarker of sensitivity or by comparing these CDK1-driven resistance mechanisms to those of the other PARPi. Niraparib resistance, for example, is predominantly driven by G1/S and G2/M cell cycle phase genes in NSCLC, whereas resistance to the other four PARPi’s also include genes from the BER and NER pathways. This difference could be due to the differing selectivity between the PARPi or their different scaffolding, as the causal contribution of CDK1 is similar for niraparib and veliparib in breast cancer and in NSCLC, with both PARPi’s having benzimidazole and indazole carboxamide as scaffolding (Slade, 2020).

**Figure 4.**
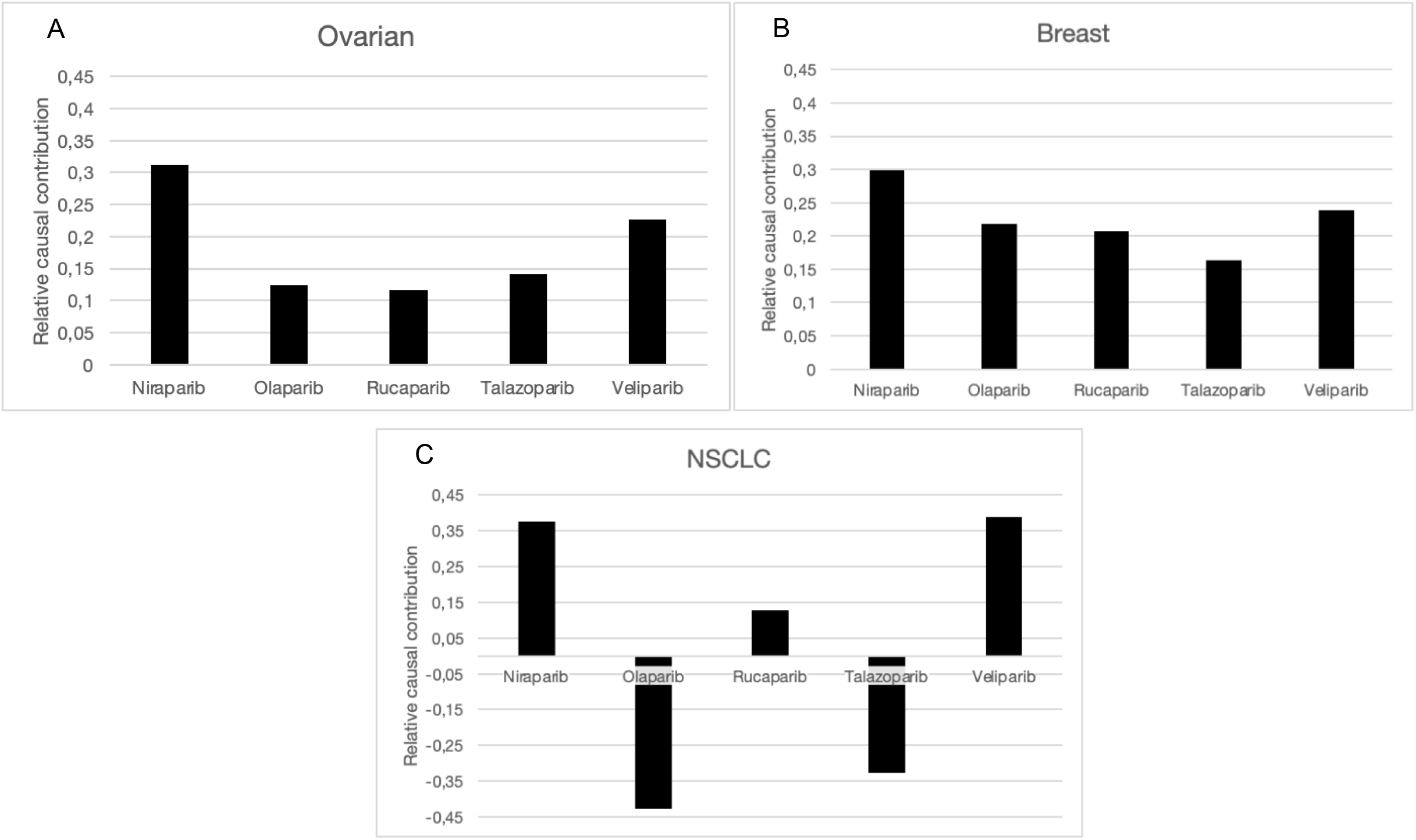
The causal contribution of CDK1 to resistance to different PARPi in a) ovarian, b) breast and c) NSCL cancer.

In addition, our analysis has also provided insight into the relative causal contributions of the pathways of genes such as ATR, WEE1 and CHEK1 to PARPi resistance, which have previously been proposed as combination partners (Figures 3 and 5). Overall, the WEE1 pathway seems to contribute the most strongly to PARPi resistance across cancers, except for olaparib and talazoparib in NSCLC where it has a potentially sensitizing effect. We also observe this effect for CDK1 in NSCLC (Figure 4C). This is an unexpected finding which can partially be explained by the observation that CDK1 plays a role in both the G2/M cell cycle phase and in HR and is known to contribute to resistance or sensitize cancer cells to DNA damage-causing agents depending on the interaction of these two pathways (Sunada *et al*., 2021). Inclusion of CDK1’s effect on HR in follow-up analyses could provide insight on why this differential effect is observed in NSCLC for olaparib and talazoparib, as WEE1 affects sensitivity to these compounds through CDK1. Additionally, olaparib and talazoparib share phthalazinone and tetrahydropyridophthalazinone as a scaffold (Slade, 2020), which may explain why only these two PARPi exhibit this effect. Lastly, CHEK1 appears to drive PARPi resistance together with the WEE1 pathway in breast cancer. Predictably, CDK1 seems to follow the same trend as WEE1 across all three cancers, with both showing their largest relative causal contribution in NSCLC for niraparib (0.38 and 0.92, respectively).

**Figure 5.**
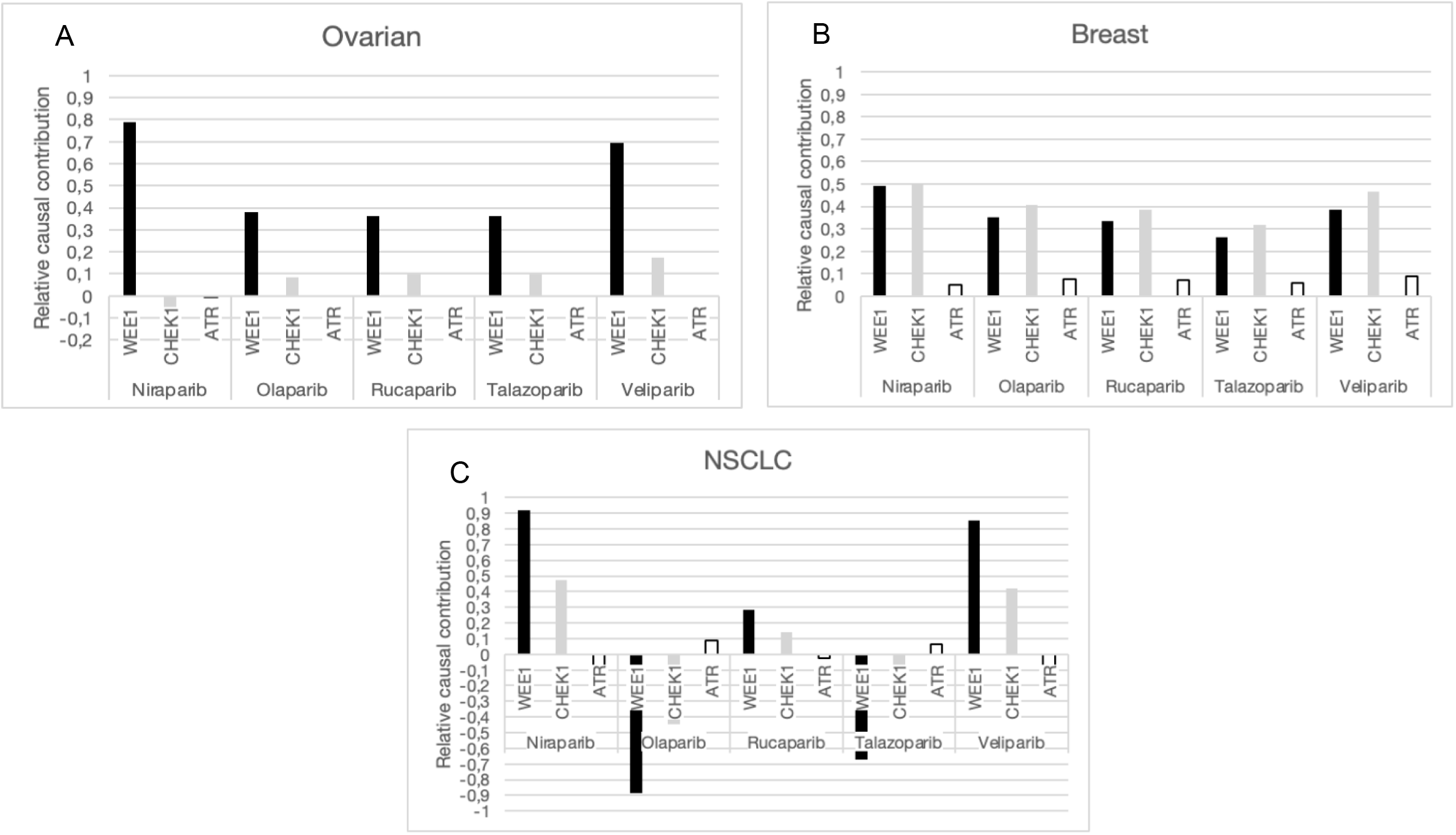
The causal contribution of the WEE1, CHEK1 and ATR pathways to PARPi resistance in a) ovarian, b) breast and c) NSCL cancer. Positive values = contributes to PARPi resistance, negative values = contributes to PARPi sensitivity.

### 3.2. Signaling pathway evaluation of CDK7 and CDK7 inhibitors such as THZ21021

Here we have focused on the DDR interplay between CDK7 and PARP inhibition and identified that CDK7 drives resistance to PARP inhibition in breast and ovarian cancer, predicting a synergistic effect of dual CDK7 and PARP inhibition. From our results, CDK7 appears to drive PARP inhibitor resistance via two different pathways: the resistance driven by CDK7 through its activation of CDK1 in the G2/M cell cycle phase is stronger than its contribution to resistance through its involvement in the TFIIH in the nucleotide excision repair (NER) pathway in both breast and ovarian cancer (Figure 6). This is particularly evident for niraparib in both breast and ovarian cancer, for veliparib only in ovarian cancer and interestingly not for talazoparib in ovarian cancer, which alludes to the strength of this synergy differing for certain PARPi. The synergistic effect by dual inhibition of CDK7 and PARP (through Olaparib administration) was shown in literature for breast and ovarian cancer (Shan *et al*., 2020), however, the specific pathway information was not known but understanding this could be exploited in the development of predictive biomarkers within the G2/M cell cycle genes. Additionally, this prediction was not observed in our results for NSCLC, which identifies which cancer types (breast and ovarian) are more likely to be sensitive to this synergy and can help with indication extension and patient stratification.

**Figure 6.**
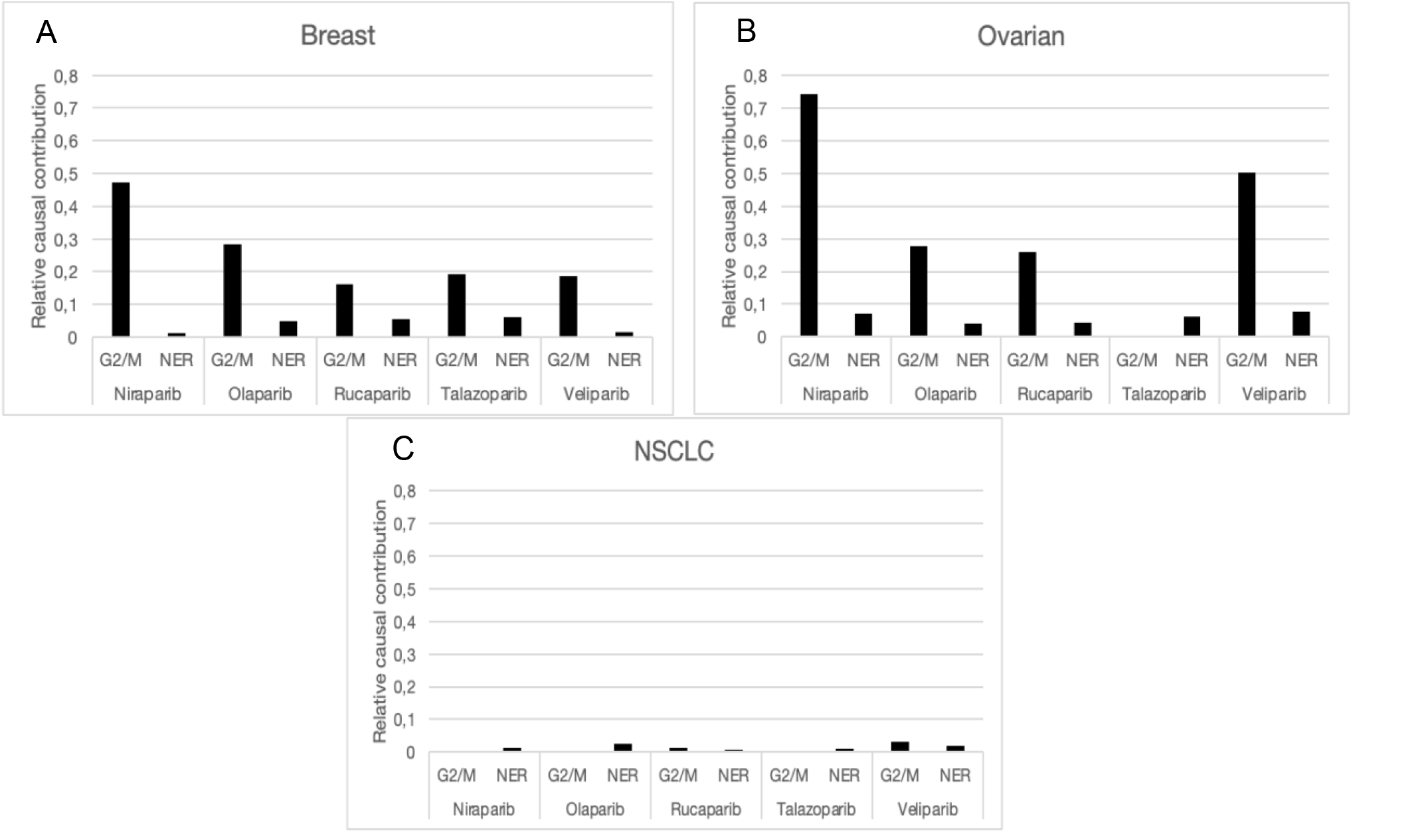
The causal contribution of CDK7’s downstream pathways in PARPi resistance in a) ovarian b) breast and c) NSCL cancer.

Our results have also revealed novel interactions specifically in the base excision repair pathway and predicts that POLG or POLG2 are significant drivers of resistance to the CDK7 inhibitor, THZ21021, in especially ovarian and breast cancer and to a lesser extent NSCLC (Figure 7 and 8). Cyclin B and other G2/M cell cycle genes also contribute to CDK7i resistance in breast cancer. CDK7 has been shown to be necessary for formation of cyclin B and CDK1 complex assembly (Larochelle *et al*., 2007), however it may be possible that CDK7 inhibition is less effective in cells that display higher levels of G2/M cell cycle progression.

**Figure 7.**
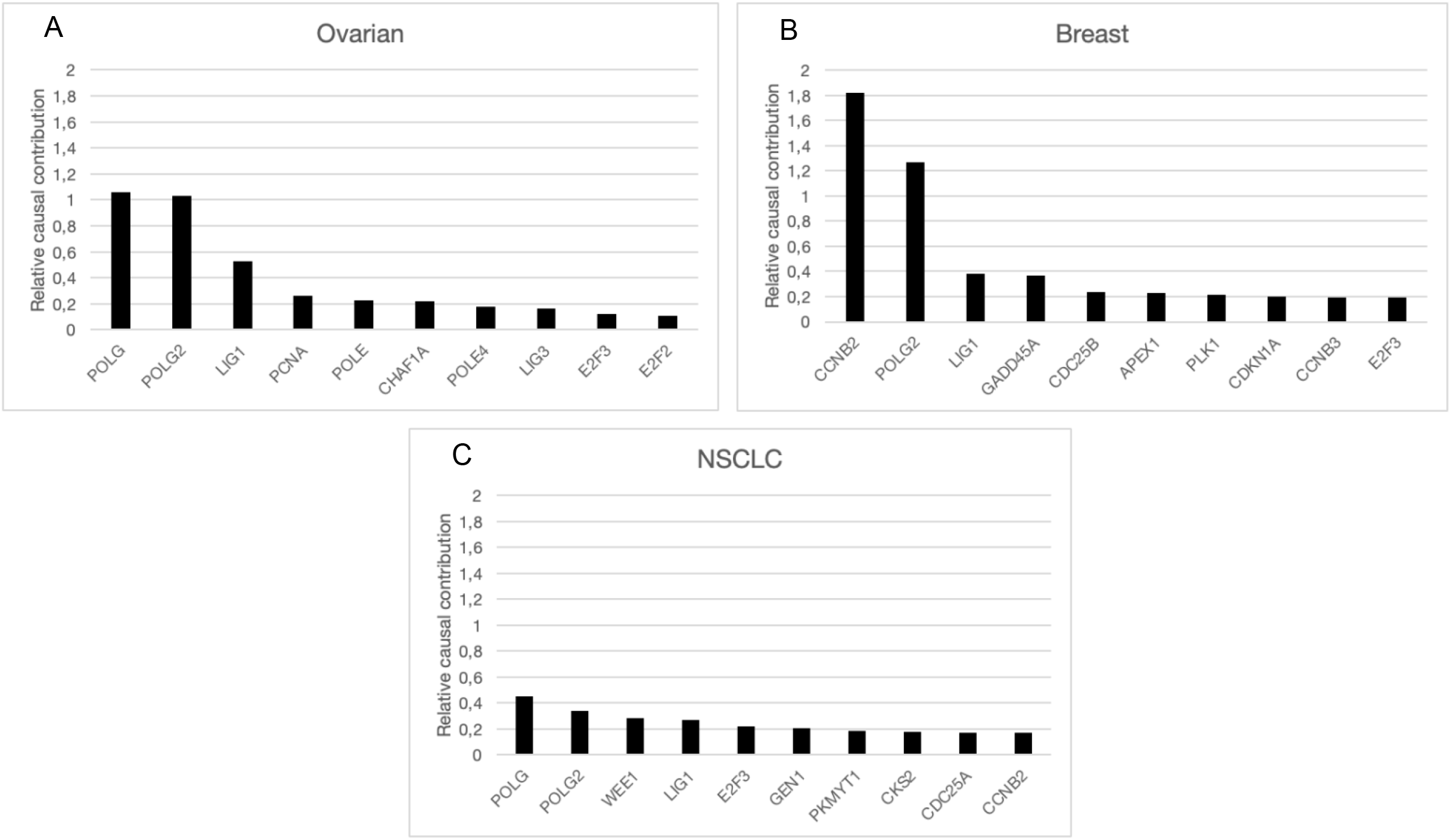
The causal contribution of the top 10 genes driving THZ21021 resistance in a) ovarian b) breast and c) NSCL cancer.

**Figure 8.**
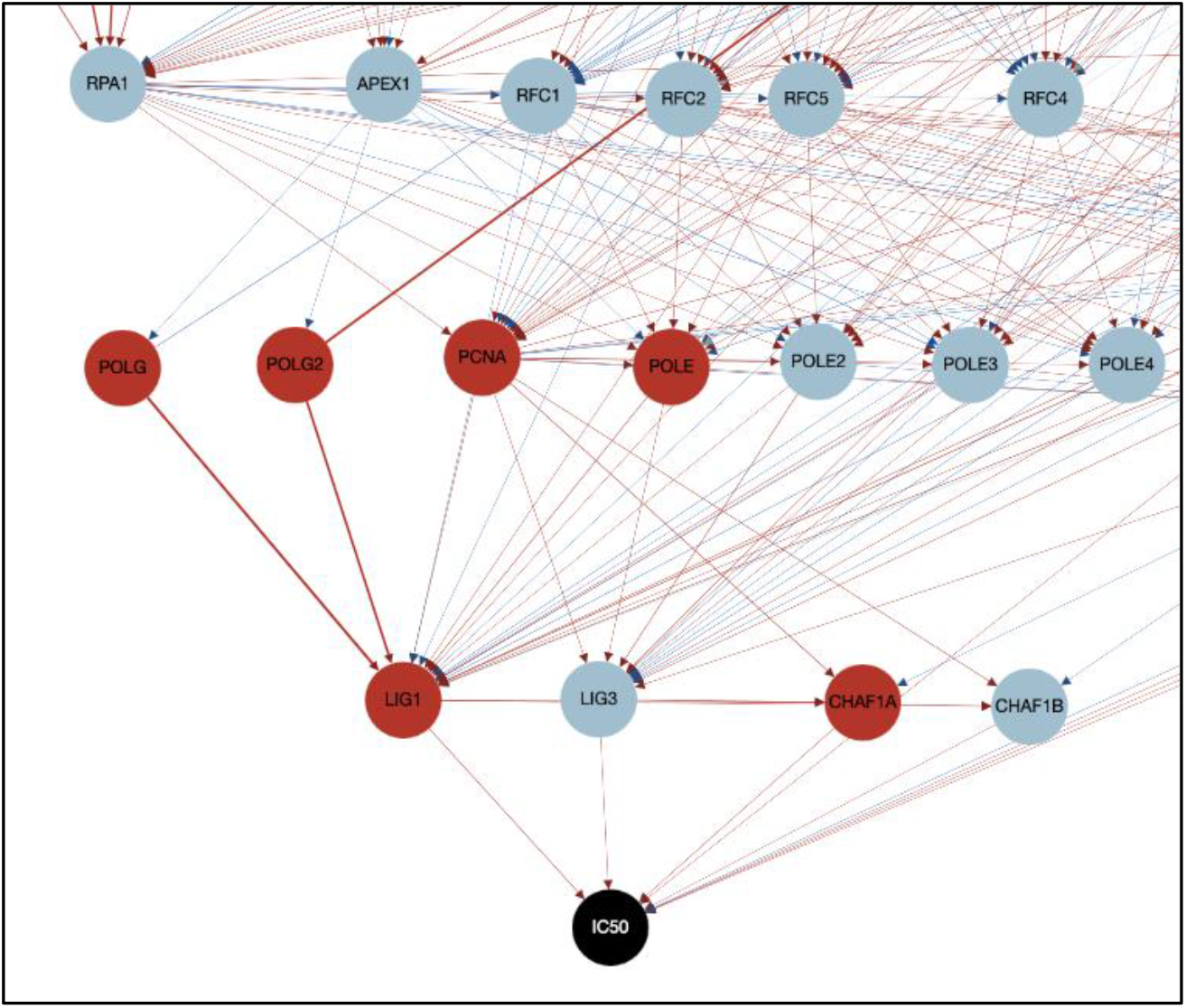
Portion of DAG indicating relation of BER genes to each other in driving THZ21021 resistance in ovarian cancer. Red circles = top 6 genes driving THZ21021 resistance.

### 3.3. Signaling pathway evaluation of ATR inhibitors such as AZD6738

Our results show that genes involved in the homologous recombination (HR) pathway (Figure 9 and 10) appear to drive ATRi resistance in breast cancer cell lines, specifically through RAD51 activation. RAD51B is the strongest predictor of resistance and appears to drive ATRi resistance mainly through the break induced repair sub-section of the HR pathway via PIF1 and to a lesser extent via the synthesis-dependent strand-annealing (SDSA)-mediated HR through the POLD’s (Figure 9 and 10). An increased sensitivity to ATRi in HR-deficient cancers in ovarian cancer cell lines has been shown in literature (Krajewska *et al*., 2015), however, the specific pathway information was not known and could be used to identify biomarkers for patient selection.

**Figure 9.**
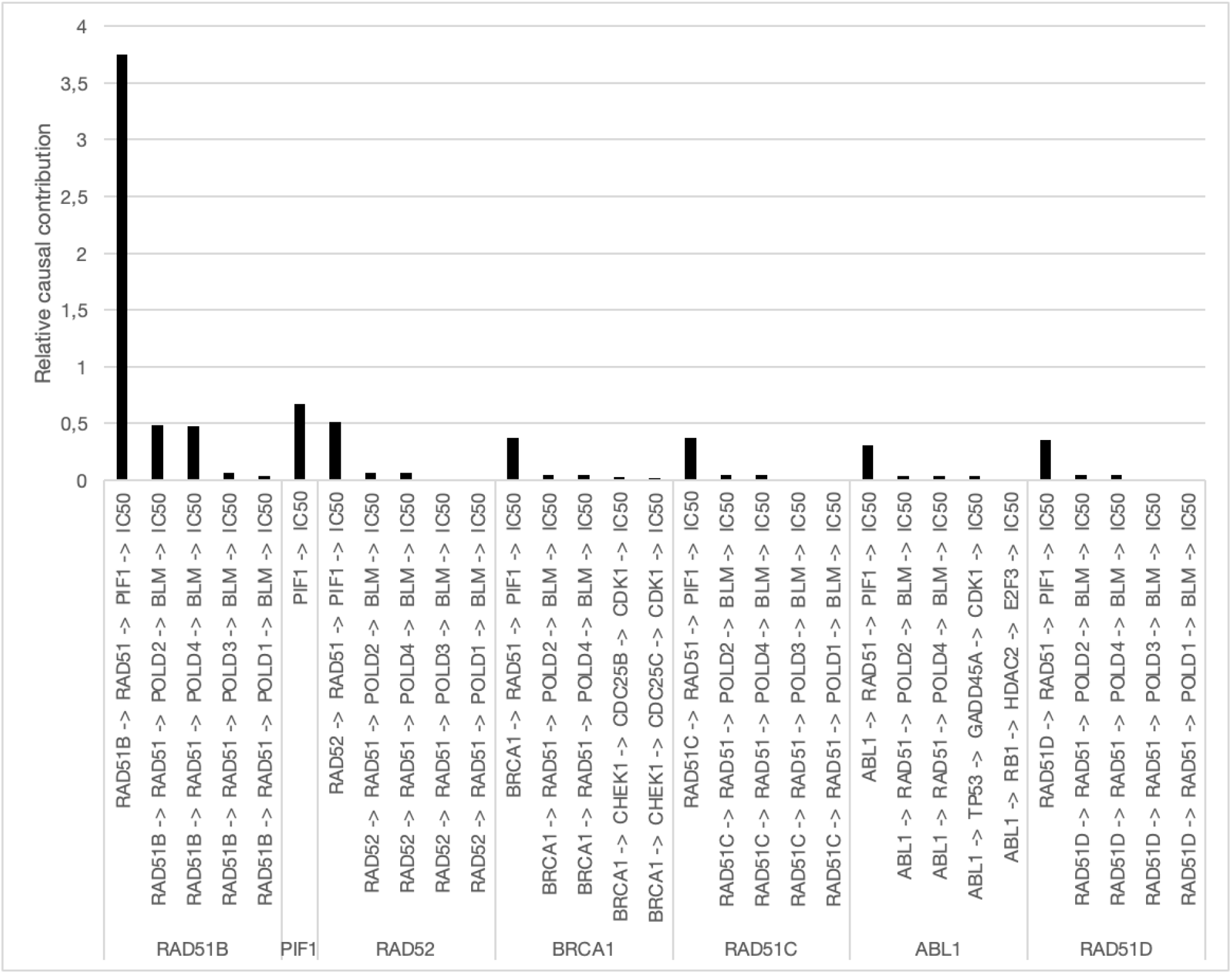
The causal contribution of the five strongest downstream pathways from the top HR genes to ATR inhibitor resistance in breast cancer.

**Figure 10.**
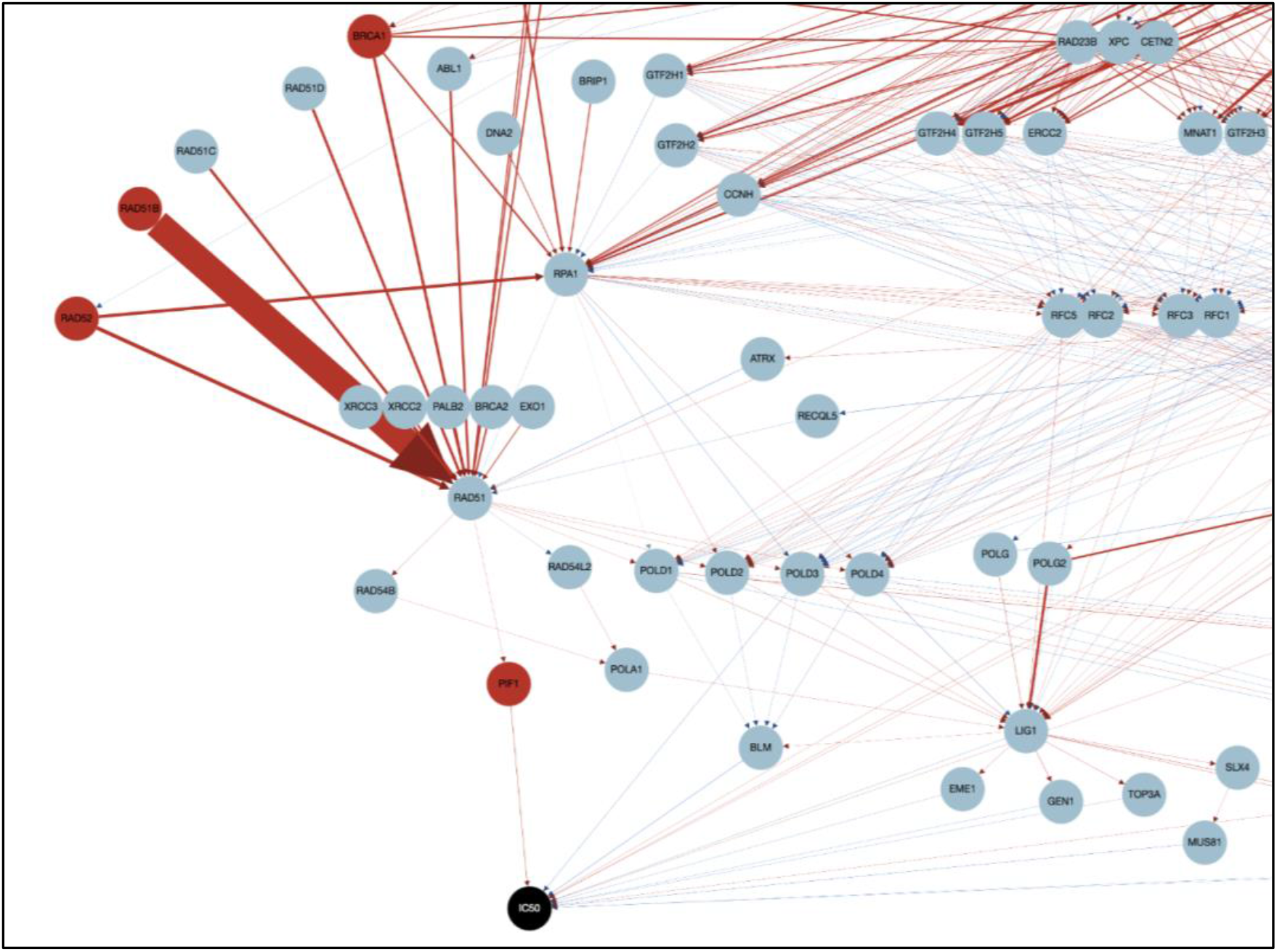
The causal contribution of downstream pathways of top HR genes to ATR inhibitor resistance in breast cancer.

Next, we compared the resistance mechanism of ATRi to irinotecan and gemcitabine in ovarian cancer to identify potential similarities or differences to these compounds, with irinotecan suggested and gemcitabine being tested as combination therapy with ATRi in platinum-resistant ovarian cancer (Konstantinopoulos *et al*., 2023; Li *et al*., 2022). Our results showed a different profile of genes driving resistance to AZD6738 compared to irinotecan in ovarian cancer (Figure 11), whereas its resistance mechanism was similar to that observed in gemcitabine (involving predominantly G1/S cell cycle genes that drive cell cycle progression through E2F transcription and to a lesser extent NER genes). This may suggest that ATRi will be more effective in combination with irinotecan rather than gemcitabine in ovarian cancer, as their resistance mechanisms differ. From our results, irinotecan resistance in ovarian cancer seems to be driven primarily by HR genes, especially RAD51D, and TFDP1 - the latter being involved in gemcitabine and AZD6738 resistance as well. The involvement of HR genes in irinotecan resistance is supported by the observation that HR-mediated repair of DNA double-strand breaks after irinotecan administration reverted cleavage-sensitive cancers to cleavage-resistant cancers, contributing to drug resistance upon re-exposure to irinotecan (Kumar *et al*., 2023).

**Figure 11.**
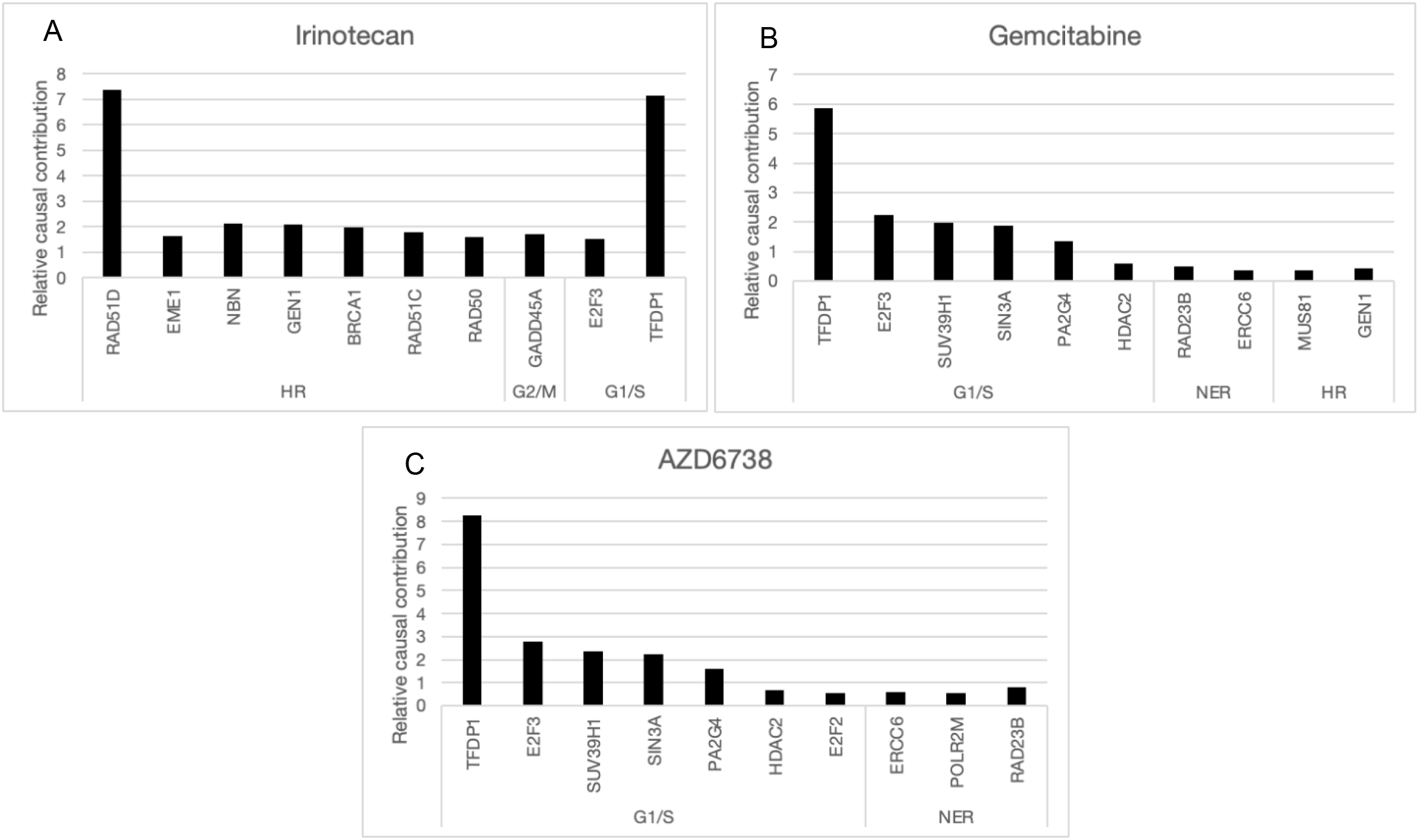
The causal contribution of the top 10 genes involved in resistance for a) irinotecan, b) gemcitabine and c) AZD6738 in ovarian cancer.

The observed differences in genes driving resistance in irinotecan and gemcitabine could be explained by their differing mechanisms of action. Irinotecan prevents re-ligation of the DNA strand by binding to the topoisomerase I-DNA complex and blocks progression of the replication fork, which induces replication arrest and lethal double-stranded breaks in DNA (Kumar & Sherman, 2023). Gemcitabine is a nucleoside analog which inhibits DNA synthesis, causing the DNA strand to be incompatible with further processing by polymerase enzymes but also rendering it resistant to excision by DNA repair mechanisms (de Sousa & Monteira, 2014). Additionally, a study in ovarian cancer cell lines found that disrupting HR through BRCA1 depletion sensitized these cell lines to topotecan (another topoisomerase inhibitor) and other agents that cause damage repaired by HR, but did not sensitize these cells to gemcitabine (Huntoon *et al*., 2013). This supports that HR may not play a significant role in gemcitabine sensitivity. In their study, ATR inhibition reduced cell viability independent of BRCA status (although BRCA depleted cells were more sensitive to inhibition of ATR), indicating that ATRi affects treatment sensitivity through pathways other than HR, which could explain why HR is part of ATRi’s resistance mechanism in breast but not ovarian cancer in our results.

## 4. Conclusion

We applied ALaSCA to evaluate the DDR signaling pathways and how their components impact resistance to PARPi, CDK7i and ATRi in breast, ovarian and NSCL cancer. We identified opportunities around these inhibitors, including the development of predictive biomarkers such as CDK1 for niraparib resistance, potential synergistic combination partners such as CDK7 and PARP in ovarian and breast cancer, and lastly patient stratification for ATRi in HR-deficient patients. Our causal pathway approach enables the generation of these insights together with *in silico* evidence for a hypothesis of how a gene’s associated pathway drives resistance to the inhibitors in each cancer.

## 5. Next steps

The results above were generated using public data from cell lines and show the potential of this approach when applied to proprietary and patient sample-derived data. Partnership with industry drug discovery groups using proprietary data to rerun the above evaluations will further refine and confirm these findings, especially with the inclusion of xenograft models or patient-derived samples which more accurately reflect the dynamics within cancer patients.

Next steps could also involve further investigation of why niraparib exhibits differences in the genes predicted to contribute to resistance (such as CDK1 and CDK7) compared to the other PARPis, as well as understanding how NSCLC is differentiated from ovarian and breast cancer. This may include the addition of further DDR genes and pathways to understand resistance mechanisms in NSCLC, as overall the contribution of DDR genes to treatment resistance in our analysis was much smaller in NSCLC than in breast and ovarian cancer.

## 6. Acknowledgements

We would like to acknowledge Dr. Paul Agapow (Director at GSK) and Dr. Dawie van Niekerk (Senior Lecturer at Stellenbosch University) for assistance with the review of this work.

## Notes

### Competing Interest Statement

The authors have declared no competing interest.

