## Supplementary figures and images for "Signaling pathway evaluation of leading ATRi, PARPi and CDK7i cancer compounds targeting the DNA Damage Response using Causal Inference"

### Supplemental Material

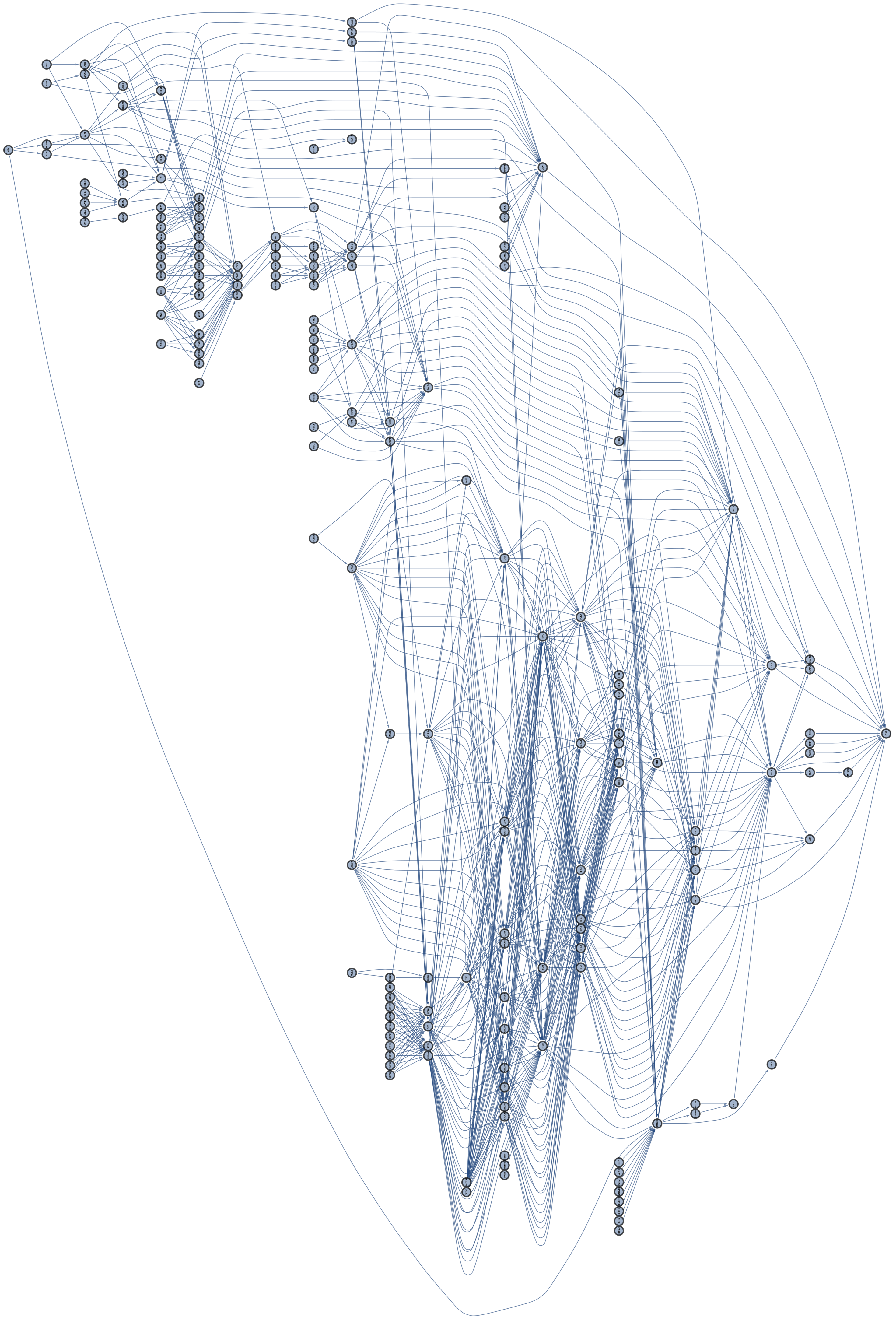
